# Association Between Relational Mobility and DNA Methylation in Oxytocin Receptor Gene: A Social Epigenetic Study

**DOI:** 10.1101/2024.08.27.609977

**Authors:** Shubing Li, Junko Yamada, Toru Ishihara, Kuniyuki Nishina, Shota Nishitani, Tetsuhiko Sasaki, Tetsuya Matsuda, Mino Inoue-Murayama, Haruto Takagishi

## Abstract

DNA methylation is a type of epigenetic modification known to exhibit fluctuations in response to environmental factors. The association of macrosocial factors, such as interpersonal mobility, on methylation has seldom been investigated. This study aimed to examine the association of relational mobility, defined as the extent to which individuals can form and replace social relationships, on the DNA methylation of oxytocin receptor genes. DNA was extracted from the buccal cells of 95 adult participants (50 men and 45 women) and subjected to microarray analysis of DNA methylation using Illumina EPIC v2.0. The findings indicate that the oxytocin receptor gene’s methylation level was higher in individuals residing in low relational mobility social environments. The CpG site associated with relational mobility is an enhancer region, indicating that social environments with low relational mobility exert a suppressive effect on the transcriptional efficiency of the oxytocin receptor gene.

**Significance Statement:** The association between DNA methylation of the oxytocin receptor gene and relational mobility was examined in 95 adults in their 20s to 60s, and found that those living in social environments with lower levels of relationship mobility had higher rates of DNA methylation of the oxytocin receptor gene. This study is a novel approach to a problem discussed in the social sciences using new analytical techniques in epigenomics.

## Introduction

Relational mobility is the amount of opportunity to form and replace social relationships and has a significant impact on human sociality (1). Previous studies have demonstrated that relational mobility is associated with a range of social tendencies and behaviors, including general trust (2), self-disclosure (3), and social anxiety (4). In social environments with low relational mobility, relationships are largely fixed and individuals are more sensitive to their negative reputations to avoid being excluded from the local community. Conversely, in social environments with high relational mobility, relationships are often fluid and open. Consequently, it is of the utmost importance that individuals adopt more positive attitudes in their interactions with others to build more desirable relationships. Although relational mobility is an important determinant of human sociality, the biological basis of this relationship remains unclear.

DNA methylation represents a valuable approach for elucidating the mechanisms underlying the relationship between social environment and sociality. DNA methylation is an epigenetic modification that regulates gene expression without altering the base sequence. DNA methylation is induced by environmental factors, and the effects of lifestyle and environmental stress have been highlighted (6). Recently, it has been demonstrated that excessive stress, such as childhood adversity (7), can promote the methylation of the oxytocin receptor gene (*OXTR*), which plays an important role in social anxiety (8). Individuals who have experienced childhood adversity exhibit a lack of trust in others (9), which may be attributed to increased social anxiety resulting from the methylation of *OXTR*. Consequently, DNA methylation may be regarded as a biological mechanism linking social environment and sociality.

This study employs DNA methylation analysis, a technique widely utilized in medical research, to investigate the relationship between social environment and sociality, a pivotal topic in the social sciences. Given the association between relational mobility and anxiety-related phenotypes (e.g., general trust, self-disclosure, and social anxiety), we focused on *OXTR*, which plays an important role in regulating social anxiety (8). The process by which relational mobility is associated with *OXTR* methylation was conceived as follows. Because of the difficulty in forming and replacing social relationships in a society with low relational mobility, individuals are compelled to adapt and conform to the expectations of others. Consequently, individuals residing in fixed and stable social environments are more susceptible to intense social stress, which, in turn, leads to increased *OXTR* methylation.

## Results and Discussion

The reliability coefficients for relational mobility were sufficiently high (Cronbach’s alpha = .89). Additionally, there was no association between relational mobility and age (*r* = -.16, *p* = .118) or sex (*t*(90.56) = 1.27, *p* = .208). A multiple regression analysis with robust standard errors revealed that relationship mobility had a negative effect on the methylation of the CpG 11 (*β* = -.325, *SE* = .095, 95% CI [-.513, -.137], *p* = .0009, Fig. 1). However, the methylation of other CpG sites was not associated with relational mobility (Table 1).

**Table 1.** Effect of relational mobility on methylation at each CpG site distant from 95% CI.

| | probe id | distant from<br>TSS | $\beta^a$ | SE | 95% CI | | p-value |
| --- | --- | --- | --- | --- | --- | --- | --- |
|  |  |  |  |  | lower limit | upper limit |  |
| CpG 1 | cg14483142 | -456 | .046 | .095 | -.142 | .234 | .627 |
| CpG 2 | cg25085537 | -439 | .005 | .069 | -.131 | .142 | .937 |
| CpG 3 | cg03710862 | -428 | .093 | .118 | -.142 | .328 | .434 |
| CpG 4 | cg17036624 | -301 | -.181 | .111 | -.401 | .039 | .106 |
| CpG 5 | cg00247334 | -243 | .042 | .074 | -.104 | .188 | .568 |
| CpG 6 | cg25140571 | -137 | .099 | .111 | -.122 | .321 | .375 |
| CpG 7 | cg23391006 | 21 | .047 | .118 | -.187 | .281 | .691 |
| CpG 8 | cg09353063 | 208 | .093 | .121 | -.147 | .333 | .443 |
| CpG 9 | cg17285225 | 296 | .121 | .108 | -.093 | .334 | .266 |
| CpG 10 | cg00078085 | 708 | -.023 | .084 | -.190 | .145 | .790 |
| CpG 11 | cg03987506 | 751 | -.325 | .095 | -.513 | -.137 | .00089 |
| CpG 12 | cg11171527 | 1094 | .167 | .133 | -.097 | .430 | .212 |
| CpG 13 | cg19619174 | 1161 | -.023 | .111 | -.244 | .197 | .833 |
| CpG 14 | cg12695586 | 1223 | -.043 | .113 | -.267 | .181 | .705 |
| CpG 15 | cg27501759 | 1585 | -.054 | .114 | -.281 | .172 | .635 |
| CpG 16 | cg04509941 | 1737 | -.050 | .096 | -.241 | .142 | .606 |
| CpG 17 | cg02192228 | 1764 | -.189 | .113 | -.414 | .035 | .098 |
| CpG 18 | cg04523291 | 1799 | .028 | .113 | -.197 | .253 | .807 |
| CpG 19 | cg15317815 | 1994 | .270 | .109 | .053 | .486 | .015 |
| CpG 20 | cg03257388 | 2087 | .298 | .111 | .077 | .519 | .009 |
| CpG 21 | cg00385883 | 3041 | -.001 | .077 | -.155 | .153 | .988 |
| CpG 22 | cg13456129 | 7514 | -.074 | .119 | -.310 | .162 | .534 |
| CpG 23 | cg04509798 | 10486 | .061 | .120 | -.178 | .299 | .615 |
| CpG 24 | cg04509785 | 11263 | -.181 | .105 | -.390 | .027 | .088 |
| CpG 25 | cg26455676 | 13841 | -.094 | .061 | -.215 | .027 | .125 |

<sup>a</sup> Standardized partial regression coefficient of relational mobility after controlling for age and sex.
TSS: transcription start site, SE: standard error, CI: confidence interval

**Figure 1.**
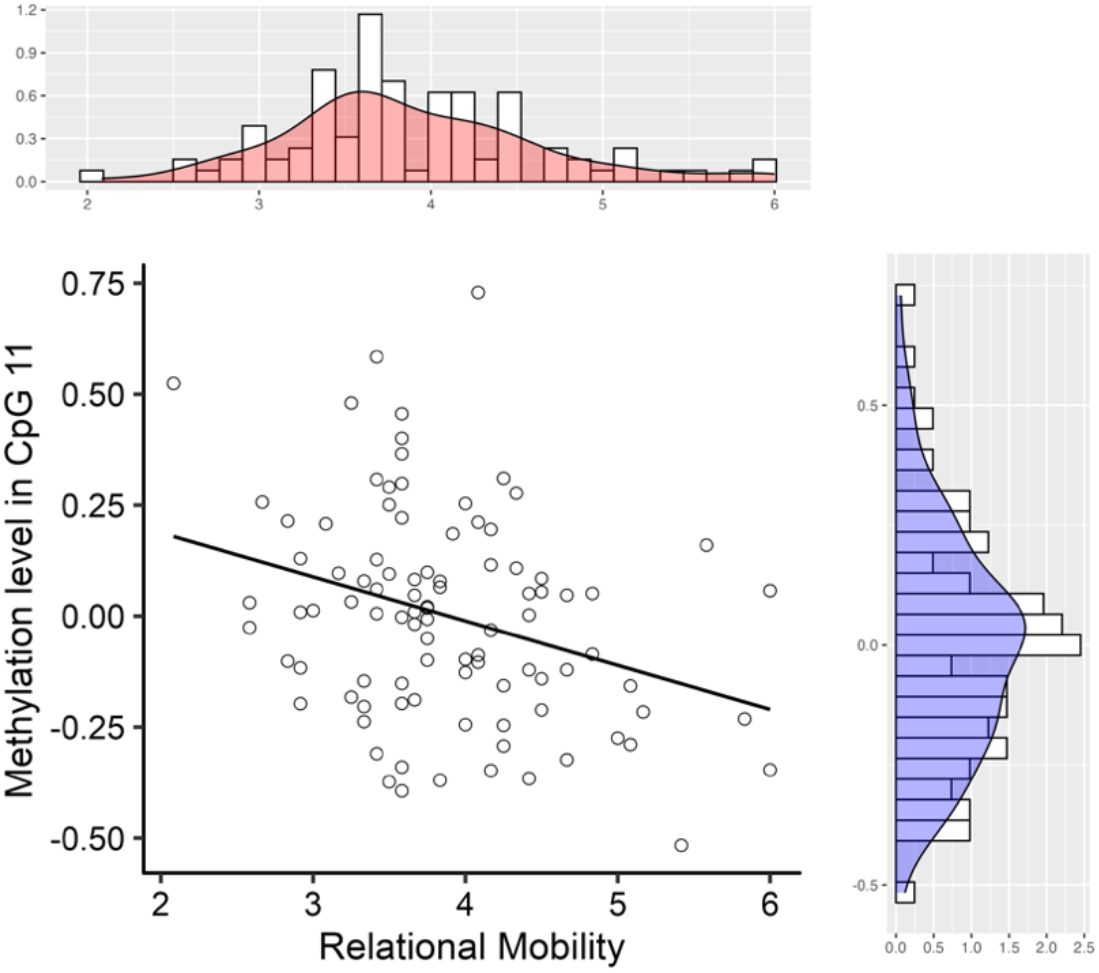
Association between relational mobility and methylation level in CpG 11. Methylation levels show residuals controlled for age, sex, and percentage of epithelial cells.

The findings of this study indicate that individuals residing in societies with high relational mobility exhibit lower levels of *OXTR* methylation, whereas those residing in societies with low relational mobility exhibit higher levels of *OXTR* methylation. These findings indicate that the expression of *OXTR* is suppressed in individuals residing in low relational mobility social environments, resulting in reduced action of the oxytocin system. Although factors such as mental illness and lifestyle have been shown to facilitate DNA methylation, it is noteworthy that the structure of social relationships can also influence DNA methylation. The findings indicate that relational mobility is not associated with the overall methylation of *OXTR*, but rather with the methylation of specific CpG sites. CpG 11 (cg03987506) is located 751 bases downstream of the transcription start site. This region functions as an enhancer, that is, sequences that increase transcription efficiency. Because methylation of this region is presumed to reduce transcription efficiency, it can be reasonably inferred that low relational mobility weakens the action of the oxytocin system.

The findings of this study indicate that DNA methylation may be a biological mechanism underlying the diverse phenotypes observed in individuals affected by relational mobility. Previous studies have demonstrated that individuals residing in social environments with low relational mobility exhibit low levels of general trust (2) and high levels of social vigilance (10). Other studies have demonstrated that polymorphisms in the *OXTR* are associated with general trust (11) and brain structures in anxiety-related regions (12). As stated previously, oxytocin attenuates social anxiety. Consequently, a reduction in the action of the oxytocin system due to DNA methylation may be associated with a reduction in general trust. Therefore, the association between relational mobility and general trust is likely mediated by the methylation of *OXTR*. In a society with low relational mobility, it is more adaptive to have a moderately high level of social anxiety to avoid a negative reputation and social exclusion. Regulation of the oxytocin system by DNA methylation may be the biological underpinning of the adaptation of individuals to the social environment after ontogeny.

The application of epigenomic analysis to social science issues can be termed social epigenomics, a methodology that has the potential to revolutionize social science research. For instance, cultural neuroscience endeavors to elucidate the genesis of cultural differences in psychological characteristics regarding the coevolution of genes and cultures (13). However, genetic polymorphisms remain static throughout life and are, thus, insufficient to explain cultural differences in the mind, which change over a relatively short period depending on the social environment. Given that DNA methylation is a dynamic indicator of gene expression that is regulated by the social environment following individual development, DNA methylation may serve as an important biological basis for cultural differences in the mind. Furthermore, given that children’s sociality is shaped by their social environment (14), it is crucial to investigate the influence of the social environment on DNA methylation and, subsequently, the development of children’s sociality. Future research should endeavor to explain cultural differences in the mind and the developmental study of sociality using DNA methylation.

## Materials and Methods

This study is a secondary analysis of data collected from previous projects (Table S1). Ninety-five adult participants (50 men and 45 women; average age = 51.4 years; standard deviation (SD) = 9.9 years; age range: 32–68 years) with relational mobility and DNA methylation data from the constructed database were analyzed. This study was approved by the Ethics Committee of the Tamagawa University (approval no. TRE23-0046) and all participants submitted a consent form before participating in the experiment. The protocol of this study was conducted in accordance with the Declaration of Helsinki. DNA was extracted from the buccal cells and microarray analysis was performed using the Illumina Infinium MethylationEPIC v2.0 chip. Based on the aforementioned hypothesis, methylation data for 25 related regions of *OXTR* were extracted from approximately 930 K genome-wide data and analyzed (Fig. 2). Because the methylation data covered a wide range of the entire *OXTR* region, each methylation data point was analyzed separately. Therefore, the analysis used p-values after correcting for multiple comparisons. Participants responded to a relationship mobility scale (12-item Likert scale). Further details of the methodology are provided in the Supporting Information.

**Figure 2.**
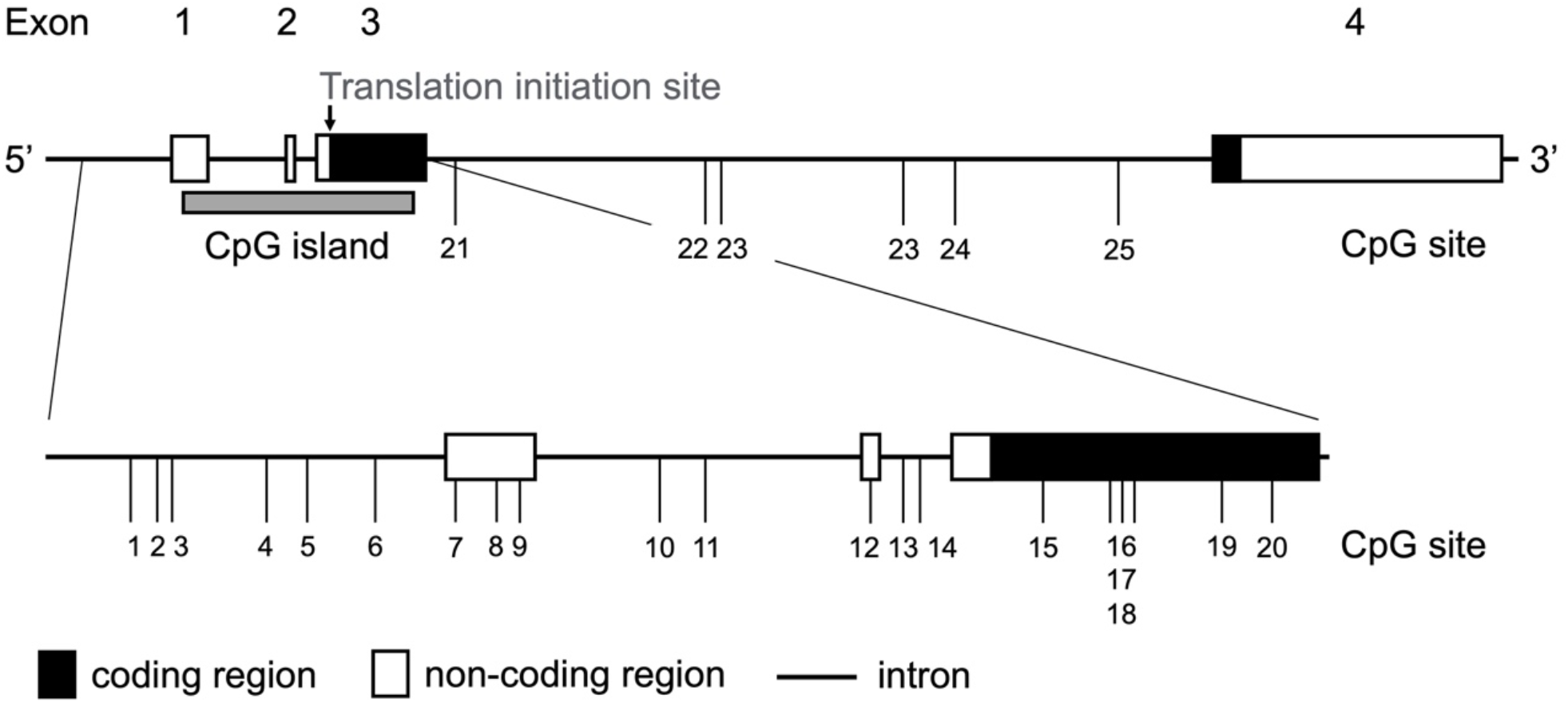
Structure of the oxytocin receptor gene. The locations of the 25 CpG sites targeted in this study are shown.

## Acknowledgments

We extend our gratitude to Yoshie Matsumoto, Yang Li, Toko Kiyonari and Toshio Yamagishi for their invaluable contributions.

## Data Availability

Behavioral and genetic data are available at https://osf.io/k87fe/?view_only=488565907a724085b95ca07234fa80da

