## Supplementary Information for "Association Between Relational Mobility and DNA Methylation in Oxytocin Receptor Gene: A Social Epigenetic Study"

**This PDF file includes:**

Supplementary Methods
Figure S1
Table S1-S26

### Supplementary Methods

#### Participants

This study is a secondary analysis of data collected from previous projects (Table S1). In this project, approximately 400 adults from the general population participated in repeated experiments, and behavioral, MRI, and genetic data were collected. Although many studies have utilized this dataset, this is the first to examine the association between relational fluidity and oxytocin receptor gene methylation. This study was approved by the Ethics Committee of the Tamagawa University (approval no. TRE23-0046) and all participants submitted a consent form before participating in the experiment. The protocol of this study was conducted in accordance with the Declaration of Helsinki.

#### DNA Collection and Analysis

DNA was extracted from participants' buccal mucosa cells collected in an experiment conducted between November 21, 2016, and March 3, 2018. The DNA was cryopreserved and mailed to Tempus, Inc., U.S., in September 2023 for commissioned microarray analysis. Microarray analysis of methylation was performed using the Infinium MethylationEPIC v2.0 chip and analyzed using the iScan system. Based on our hypothesis, we used gene annotation and promoters (GENCODEv41; https://zwdzwd.github.io/InfiniumAnnotation) to extract data on the CpG sites corresponding to oxytocin receptor genes from the approximately 930,000 epigenome-wide datasets obtained in this study. Twenty-five CpG sites, with the gene name *OXTR* in the annotation, were included in the analysis (Fig. S1). As the DNA was derived from buccal mucosa cells, the percentage of epithelial cells was calculated using EpiDISH (1) and used as a control variable in the analysis.

#### Measuring Relational Mobility

Relational mobility is an indicator of the likelihood of meeting new people and the ease of forming new relationships and dissolving existing ones. The 12-item scale developed by Professor Masaki Yuki was used to assess relational mobility. The content of each item can be found on the website (http://relationalmobility.org/the-relational-mobility-scale). The mean of the 12 items on the Relationship Mobility Scale was used in the analysis.

#### Statistical Analysis

Multiple regression analyses were performed for each CpG site to examine the association between *OXTR* methylation and relational mobility. The relationships between mobility, age, sex, and percentage of epithelial cells were the explanatory variables, and the methylation rate at each CpG site was the objective variable. Robust standard errors were used in the analysis. As 25 multiple regression analyses were performed, the p-values were Bonferroni-corrected ($p_{corrected}$ = .002).

>NC_000003.12:c8770071-8755774 Homo sapiens chromosome 3, GRCh38.p14 Primary Assembly

CGACAAGGAGGCAGAAACGGCTCTTGGGCGCAGACAAGCAGAATCACTTTAAATGAAGACAGTGTTGTGC
TTCAGAATTTCCTCTAAAACTACCGAAAAAATAACGCCTCTCCCAGCACTGCTTAGAATAGAGGCCATTT
CTAATTCCTCATTAACGGGAATAGGAACAAAAGTATTCCAAAGCAAAGACTTATTTGAGTTCACTGCTAA
AGCCGCTACATCAAGCTGGAGGTGTGGGGGGAGAGAAAAGCCTGAAAATTAACATCATTTTTGGGAAATA
ATCAGTTTAAATGCTTTTGTAACTTCATCACTATCTACCCGGGGAAGAACATTATTATTCAAGCCTCCTA
TGTGTCTCGGAGTCAAGAGCTTCTAAACCAAGAAAGGAAGAAACGGGCGGGTTATTGACGAGTTCCCTCC
CTCTCGCAGTTTTAAACCACTGCAAAATAAACCCATTTGTTAAGGCTCTGGGACCAACGCTGGGCGAACC
AGCTCCGCTCCGGAGGGGTCTGCGCGGCTGGCCTCGCCCGCCCCCTAGCGGACCCGTGCGATAGTGCAGC
CTCAGCCCCAGCGCACAGCGCCGCATCCAGACGCTGTCCGCGCGCGCAGCCTGGGAGGCGCTCCTCGCTC
GCCTCCTGTACCCATCCAGCGACCAGCCAGGCTGCGGCGAGGGGATTCCAACCGAGGCTCCAGTGAGAGA
CCTCAGCTTAGCATCACATTAGGTGCAGCCGGCAGGCCATCCCAACTCGGGCCGGGAGCGCACGCGTCAC
TGGGGCCGTCAGTCGCCGTGCAACTTCCCCGGGGGGAGTCAACTTTAGGTTCGCCTGCGGACTCGGTGCA
GGTAGCTGGGTGCTAAGCAGGGGTGGACGGGATGGCTAGGGCCGGTGGAGCCATCGGGACCCGAGTGGAG
GTGGTGGGGTGCCTCGCACTCCTTGTTCCTGGAGGAGCTCGGGGTGTTCCGAGAGATTGTAAAGTGACTT
CTCGGGATTGAGACTCAGAGTCCTTGATTATCTGGGTCCAAAGCGCAAGTCAGGGGTTCCAGAACTTTCG
AGGCTGCCGGGCTGGGGAGGAGCCCCGCGGGAGGTGTCTATGCCAGGGTCTGGGAACAGCGCTTGGGCAT
CTTGGGCTTTGAGGCAGGGGTTCCCCCAGCAGGGACTGCAGAACCGGGTTTCCACCGAAGCAGGTGCTGT
GGAAGTTCAGGGAGTGACGGCCGACCGTCTTAGAAAAGGGGGTTAGACGGGGAAGGACCAGAGCTGGGGT
TTCCCAGGCAAGTGGCTATTTGGGGATTTCCGGAGGAAGTACTTGAATTAATGTTTACTAGGAGAGGGGC
TGGTTTGGGGGTCCCGCGGCAGGTGGATATGCTGAGGGTCGGAGCCTGGGGCGAGTAGGTAGTTTGGAGA
TTCCCTCGGGGAGGTGATTTGGTTTTAGATTCCCACTCCCGGAGGAACGTTGCTGATTTTGACCCTCCCT
TCTCCCCCAGTGGAAGCCGCTGAACATCCCGAGGAACTGGCACGCTGGGGGCTCTGGGCTTGTGGCCGGT
AGAGGATTCCCGCTCATTTGCAGTGGCTCAGAGGAGGTACCTCCAACGGGGATTTCTGGGGTGGCGGCTG
AGCAACCCGAGGCCGGCGGTTGTGCCCTGTTGTTTCAGATGAGTTGGGTTCCTGGGAATGGGACAAGCAC
GCCCTACCCGCGTCGGAAGAGAAACGCGGCGGTCTCCTCACGGCCCTCCCGGTTTGTTTCAGGGTGGACC
CAGCAGATCCGTCCGTGGAGTCTCCAGGAGTGGAGCCCCGGGCGCCCCTACACCCTCCGACACGCCGGAT
CCGGCCCAGCCGCGCCAAGCCGTAAAGGGCTCGAAGGCCGGGGCGCACCGCTGCCGCCAGGGTCATGGAG
GGCGCGCTCGCAGCCAACTGGAGCGCCGAGGCAGCCAACGCCAGCGCCGCGCCGCCGGGGGCCGAGGGCA
ACCGCACCGCCGGACCCCCGCGGCGCAACGAGGCCCTGGCGCGCGTGGAGGTGGCGGTGCTGTGTCTCAT
CCTGCTCCTGGCGCTGAGCGGGAACGCGTGTGTGCTGCTGGCGCTGCGCACCACACGCCAGAAGCACTCG
CGCCTCTTCTTCTTCATGAAGCACCTAAGCATCGCCGACCTGGTGGTGGCAGTGTTTCAGGTGCTGCCGC
AGTTGCTGTGGGACATCACCTTCCGCTTCTACGGGCCCGACCTGCTGTGCCGCCTGGTCAAGTACTTGCA
GGTGGTGGGCATGTTCGCCTCCACCTACCTGCTGCTGCTCATGTCCCTGGACCGCTGCCTGGCCATCTGC
CAGCCGCTGCGCTCGCTGCGCCGCCGCACCGACCGCCTGGCAGTGCTCGCCACGTGGCTCGGCTGCCTGG
TGGCCAGCGCGCCGCAGGTGCACATCTTCTCTCTGCGCGAGGTGGCTGACGGCGTCTTCGACTGCTGGGC
CGTCTTCATCCAGCCCTGGGGACCCAAGGCCTACATCACATGGATCACGCTAGCTGTCTACATCGTGCCG
GTCATCGTGCTCGCTGCCTGCTACGGCCTTATCAGCTTCAAGATCTGGCAGAACTTGCGGCTCAAGACCG
CTGCAGCGGCGGCGGCCGAGGCGCCAGAGGGCGCGGCGGCTGGCGATGGGGGGCGCGTGGCCCTGGCGCG
TGTCAGCAGCGTCAAGCTCATCTCCAAGGCCAAGATCCGCACGGTCAAGATGACTTTCATCATCGTGCTG
GCCTTCATCGTGTGCTGGACGCCTTTCTTCTTCGTGCAGATGTGGAGCGTCTGGGATGCCAACGCGCCCA
AGGAAGGTAGCCAGGGCTGGGAGACCCAGGAGGAGGGAGCCTGGTGGCTGGGGGAGGCCCTTATCTTGCT
GCCTCAGAATGTCCAGGGGTCTGTGGACTTCCTGGGGGATAAGCGGGTTTGAAATCCCACAGAGTCACTG
CTCTGTCATCCCTTGACCAAGTGACTTAGGGCAATTAACCTCCCTGAGCCTCCACTTTCTCTTCTGTAAG
GTGGCAATAAGGATAAAAGTACCAACTGTCACCAGGCATAGGGGAATGCCACGAGAAAATGCAGTTAAAG
TCCTTAGCACAGTCCTTGGGCTGCATATGGGCTGTATGGTTTACTGTGGTGGTGGAAACAGGTTCAAGGG
ACTCCATCTGCTTTCCCACGTGGTTAGGAGGAGGTGGTGGAGGTAAGTTTGGAACCCCTGGCCAAGCTCA
AGCTCCTTCAACTTTAAGTTCACGTTAAGATGAACTTCCACTTTAAGTTCAAGAAATCCAGCTGAAGCCA
AGAAGTCTGGTTTGGACAAGGACAGCCTTGCAGGAGTGGCAATTTGTCCAACCAAGCACCTAGTAGTTGA
AGGGGGTGTTGGGGGGCAGAGGAGTCCAAGGGAGAGGTGAAGACAAATCCCTGAACTCCTCATCGAGTGG
AGGTGATGAGTCTCCATGCAGAGAGGTGAGTACTGCAAGGAGTGGTGGGCAGGCTTAGGGAGGAGAGCGC
CCAGGACTGAGACTTCACTTCCACTTGGAGGAAAGAGAAGAAAGCCTTGAGGGGGACATTCAAGTTTGAG
GGGGAGGCTGGGTGATTTCTGTAGGTGGGGAATGTGCCTTCCAGGTAGGAGGACTGCTGAGGACAAGGCT
GAAGGTGGGGCAATTGTCCATTCTTGGCCTGTGCCAGAAAGCTAAGAGGAGGGCATAGAAGAGGGCGACC
TAGGGAGAAAGCTTGGAGGAACAGAGGCACCCTAGGGCTCTGGGGTTTGTGGCTGGTGCTGGGTGGCTGT

CCCCAAGGCACAGAGCACCAGGCAGCTTCCTGTCACTCCCTGCCCCCCAACCCCTTTCCAGTAGCTCATT
TGGAAAGACCAGGAGGGGGTGATTGCTGGTGGTGAATGATTTATAAGTTTTTGTTTGAAGGCAATCCCAT
AACGCGTGCTGGATGCTGGGGACTTCAAATGTTTGTAAAGAAAGACCAAAGGATTTGCTTTCACCCTTTC
TGCACTCCAGACACTGCAAATGTCCCATTTCTGTGTGCAGTTACTACTGCTCCCAATAGTCTCAAATACA
TGCATCCTGGGAAGCCGAAATCTCAGTGCCAGCTGCTTTCATTCAGTTTGGTAACTTTGCTTTGGTCCTT
TCTGAAAATCACAGACTCCAGGCTTTATGGAACCCTCCCAGTTAGCTCAGTGTTTGCTTTAGTGGCCAAG
ACACTGGTTATTCAATATTTACCGAACACCTGTGGCCCCTGGTAACAACTTACAGAGAGGGTTCTAGGGC
CCCGAGAGGGGAAGGCACTTGCACCAGGTCACACAGCAAGTCAGAGGCACAGCTGACGTTTCCTATGCCA
GGGTTCTTATGAAGGACACATGATGAGGGAGTGGGGTTGGGCAAAAGCCAGTAGTGGAAGTCTCTAAGTA
AGGCCTTAGGCCTCACCTGTTTCCTGGGCAGATCTGAAGAAACCAGATCCAGGTCTTGAGGAGCTCGAAC
ACCAACTATCAAGAAAGTGAAACCAGCACCCTTTGGAGTAGATACTGGAGAGATGTGGGGTCTAGTCCTC
CTCCAGCCCTTACTAGCTTTGGGATGTTAGATGAGAGATCTCTTTATTCCCCTTGAGCCTTAGTCTTGCT
TTTAGAATGAGGACATTTCAGCATTCAGTTAATTCATTCAACAGACATTCCACTCCTCTCTCCTTGGGGT
CCCCTCGCCTCTCCCATTCATGTCTTGAGTCTCTGGTGATTTTTCTCCTTCCCTTTCCTCCCAGCTCCCT
CCTCTATGGCTTCCTGCTTTTCAGCCCCCTCACTTGCACTCCTCTTCTCACTGTGTGTCCTCCCTAAAGA
AGGAGAGTGTGTGGTCAACACGTCACACGTACCGTGAAGCGAAGTTTAGATCTTTTCTACACTTTAAAAG
CACGTCAACAATTGATTTCTTTGTCTTAGAAGCAAAAGAGAGAGGAAAAAACAGACACACACAGATCCTG
GCCAGAGTGGAGGTCATAGGCCTCTTGAGTGTTTGTGGTCCTCTGAGGATTTTCCTGGGGACCAGACCTT
CAGTAGAGACGGGCTCCTGCTGAAGCCCAAGGGGCAGGGAAAGAAGAGGACTTTAAGAATTTCAGGTCCT
ACTTCCCTTTGCTTCTTTAGGAATTCACAGGGAAGGGTGGGTGTTGCATGAAAAACCCAGGGCTTCTCTT
GGTCCTCCTCAGCATTCTGTCTCAGGTACTGAGAAGAATCCAAACACAGACCGGCAGGGATTCATGATGC
TGCTGCAGCCGGGACCTCCCCGCTGCCCCAAGCACACAGCCAGGAGCAGGAGCCCAGTCCAGATGTGAGC
ACCTCATCAAACACAGAGGTGATGACGTGGACTCTGGCCACCACCAGCCTGGCCACATCATGCCTAGGCC
TCGGGGATGCAGGGGTCAGAGTGGCTGTGAGCTCAGTTAGAACACCAGCCGTGTGTCCACTTGGCCTTAA
TTTGGATAATTGCAACTGGCAGGTGGAAATGGTGTCTGCATCACTTTGGAGATACCTCTCAATCCAGAGG
CTACCAGCAAGTGTTAAGGAAGGAATAAAAGAGAACCACACTTGAGCTACTCAAGAGGTTATTATCGGAA
AAGTGGAATTAATGCACAGCCCCACTTTCCTCTCCAAGAGGACAGAATTTAGATGCGGAAATATCTAGAC
CAGGGCCCCTTACTCCACTCATCACCTGAGGAGCCAGTACTGAAGCCTGTGGCTAACAGTGTGATGCCAT
CCAGGGATGCTTGCAGGTGGAGCAGGCTCACAGGGAGGCAGGAGAGGTGGGGCACACATGGGTCTGTTCC
TGCTTTATCATGAAAACCCCAGGGAGGTAGGGGAGCTTAGCCAATGGCCACCCATCAAGGCTGCCAGAGT
GGGTGGGGAATGTCACGCTTCTGGTGGCTGTAGATGTGGTGGCCCAGGGACCCGAAGTCCCTCTCTGCTT
CAGCATTTGCTCTCTCTTTTCCATCTCTCCCTTGAGCAACTTCATTCCCTCATATGCCTTCACAAATACC
AAGGGCTGCCTTTGTGGGTTTAAAGCTCACCTTGTTCCAAAAGTAATGTGCAGTGGTGGCCAAAAATTTA
GGTGCCTCAACAAGATAAAATAGATATAAGGAAACTGGGATAAAGCAGGCAGTAGGGAGAAGAAAAGGAA
AAAAAACTATTACAATAATAGTAATAATACTATTAATAATAATAGTCTCTCTGTGTATAGACAAGCTCTA
CACCTGGTTAGTTGTTATAAGAACCAAAGAGGAAAACTTTAAGTTTAAATATTTCCATGATTCCGAAGAA
AGAGCCAATGGGAGCATCTGGGTAATTTGATGAGATCCATTCTTCAAAGCAGCCGGATGCCGTCGGGTGC
TGACTTGGTGAGGCTGGTGAGACTGGATGCCCCATGCAGGGTTTGACAAGCCCTTTATATATCTTTCAGA
ATTAAATTCCAAAGCCAAAGATTCGGCACCAGCCTTTTCCCTTGAACCCAGTGGCATTTTCTATGCTGAT
TGCAGGGGCTCTGACCCTGGGGCAGTTCTGAGGGAGGAGATGAGGATGAGGAGGGAGTTTGGTAGAGTGG
GGCTGTTTGCATTTCTAGGGTTGGTTGATGTTGATTGGGTGCGGTGGACAGGATTTGGGGTCTCTGTCTC
TAGTCTTCTCCTTCCAGCCTCTCCCTGGTAAAACGCCCAGTCCAGTGTCCTTAAGCCAGTTGCTTTACCT
GCTCTGAGCCTCCACCTCCTCCTCCATGTAATAAGGATAGCACGGCTCACTGCCTGGGGTTGATGTGAGG
ATTAAATGGTGTGACCTATGGCCCCTTCCTGATGGACAAGATGGGGATTCTTGTCTTGAGCCTGGGTGGA
GGGAATGGGAGTGGACAGCAGGTGATGTGTCCGGGGAGGCTGGGAGGTGGGAGAGAAGCTTGTTAAGGCC
GGAGAGGTGGACTATGAGCCCTATCACAGCAGGGACGCTGCTTGTCCTGTTCCCTACTTGTTAAGCCCAA
GTCCAATGTGTGGGCTGTTTTGTGGCACAACTCCAGGGGATGCCTTTTACCCCAGGGGATGCCTGGCACA
AGGCTGCACAGTCATTGATAGCTCAACCAGTGCTTGCCAAATACGTGGAAAATGGGAGCAGAGAGGTGAG
TGAGACGGGGCAGGGATGGCCATGAAGAGCTGGACTCAGGAGGAATAGGGACTTTCTAAGCAGCTGTCTG
AAACAGAACTGGCAACCTGGAAAAGAAAGGAGCGCCCCATCACTGGGGCAACCAAACATCTGTCAGGAGC
GTTGGACTGGGAGGAAGGCTGACAGGCACAGGAGGCCTGGTTTGAACTGTTTCCCCATCTGTAGAATGAG
CTTCCCAGCCCCTCCCTGTTTCTGTGGGACTGAGGATCCTCGGGCATGTCCCGTACACCTTTCCTTTCCA
TGAATGCTGTGCCCGAGGTGTGGGGGACCAGGCCCCTTGGAGGGGCTTTTTCTTGAGGTGACCCAGTGAT
GTGGAGAGCATGGTGGGCCGGCAGGAGAAGCTTGGACAGGATATTCACTCTGGGCCATGAAGAGTGCACA
AGCTCGGCCCTTCTGGGTACAAATCACAATTAGATGGACAGTTAACTTCAGACACTTATATGATTAAGTA
ATTGTGTGTTCTTTTATGTTCTTAGCTAATGTTAATGTTTTCCCTTAGGGTTTCTGCCCTGTGGTGATTT
CACTTCTTTTCCCCCCAAAGGTTTGATAAATAACAGCTAAAAAGACAAAAATCCAAAATCAAAAAACTCA
AAAGCTAACCCCAAACCAGCCTCTTGGGTGAGGCCCTGCTACATCATGTTTCCATTTGATGGCTGTCTGT

GTGGGTTTCGGGAGGGGTGAGGAGGGGAAGAGGGGTTGTGGAGGGCCTCGGATCAGGAGGAGATTAGCAG
GTGCCCTGGCTGCATCACACTGTCTGTGGAGGCTGGGTGAGGTCTGTGGCCCCGGGCTGGCGTGTCTGGG
TGTGACCTGAGGCACTGGCCCACCTGCTGACCTTACCACCTTGCCCTGGCAACTTTTGGAAGCAAAGGCA
ACCCCACAGGGCTCCCTCCCAGGGGCTGAAGCCGGACCCTGTACACCTGAGCCCACCTGCCAGAGCCCCC
AGCCCCAGGAGACAGTAGTGGGAAGTAGCCAGTCCCCTTCTGGGACCTGGTCTTGTTGCCATCTGTGTCA
TTTGGGGCAATGTCTCTTGATGGCCTCTGGACCTCTGCCATTCCCATCTGAGCAATGGGGATAACGGCTC
ATAATGGAGCTTTGACACCCTCTCTGGACCTCAGCTTTGGTGAAGGGAGGAGTGGGATCAGTTGGTAACA
GCAGCCACCACTGGCTGAGTGCCCACTGTGAGCAAGGCCTGTGCCCAGGCCATTTCTGCTCCTCACCCTA
ATGCAGCAAGGAAGGGGCCCATCATGGCTTTTCTACAGATGATGAAACTGAGGCTTAGAGAGGTGTGACC
TGCCCACGATCACTGCTGAGTAAGTGATGGAGCTGGGATTGGAACCCAGCCATCTTACACTAGAGACTGA
CACACTGCATGACGACCCCAACCCCTGCCGCCCCAGGCCAGCAGTTTTCCCGATGATCTACACCTTCGAC
TTGGGGATCTCCTGGGCGCTTGTTAGATCGCTGGACCTGCCTCAGATCCATGGAAGCTGAACCCTCCCCC
ACTATCTGGCCCCAAGGTCCTGACATCTTGGGCTCTTCCTGCAACACCTACACACACACACACCGAGGTA
CACATCCGCTGATGGCACTGCCCTCCCAGCATTCTGGGTTCAGTGCCATGCCCCTCTCATCCTCCCTGTG
TCCACCAGTTCTATAATCTCAGCTGTACCACATGCCTCAGGTTGGCACTTTGTTTCTCTTTCCATTTTCA
CCTTCTTGGTTTAGAGCACGATTTTTTCAACTTTGGCACTATTGACATTTTAGGCTGAACAGTCTTTGGC
GTGTGTGTATGTGTGTCAGGGAGGAGCTGTTCTGTGCATGGTAGGATATTTAACAGCATTCCCGTCCTCC
ACGTGCTACGTGCCATTAGCAACCTCCCTGCCAGCTGTGGCCACCAAAACTGTCTCCAGACATTGCCAAA
AGTTCACTAAGGGGCAAAGTCATCCCTGGTTAAGAACTGCTGGCTTAGACCCCTTGTTCTGCTTTTTAAA
CTAATACCTGGCAGTGAGTACTCTCTTGTTGCCCTGGTCTCTGGCCAGCCTGCCTTGTGTACCCCCAGCC
AGGCCCGATGATTTGCCGCTTTCCACAAGTTCCTATGGCCTTGCTGTCTTTGCACCTTTGCTAATACTGT
TCCTGCCATGGGAACGCCCACCCCAGTTTCTTCCACGGATTCCCATCCTTTGAAGCCCAAGTCCATTGTG
TGGGCTGCTTTGTGGCACAACTCCAGGGGATGCCTTTTACCTCATGGTCTATGAACAGGGTGTTCCTAGG
AGCTTTGCAGTGCACAGCCTGAACAGCTGAACATGGCAGCCTCATCCAGTGCCCCTTTCAGGAAACCATC
CCTGTTTTCCCAGTTTGCGGGGCTTCTTCCTCTGAATGCTAAGTCATTCACTCCAAAAAACAGTTACTGA
TGCCTCTGCAGGCTCCTGTCCACACAACCTCTGCTTTCATCTGTCAACCCCCAAGTGAAGAGAGTTGGCA
AGGGGCCCTCCCAAGCCCTCGCCCTCCTGCGGCCAGCTGCTGACATGCATAAATGTGGGACCGTGTGGTC
AAGGTTGATCACAGAGTTCTGGGTACACCCACAGACCTCTGGGAGGGACACCCACAGTTTCTGCTGGGTT
AGTGTCAGATGGAGCGGCAGTGCTGACTTCGGGTGTGGGGCTGAGCCTGCCCACAACTCAGGTTTCTGGG
GTTCAGAATCTCAGGTTTCTGAAGTTCAGAATAGGACTTAGGTTGTATACAGGGTGGAATATCCTGCTGG
TGCAAGTCAGTGTAGCTGATGGAGTGGGAGTCATTTTGGGTACAGGCCTAGCATTTTAGCAAAGGGGCTG
CAGGTCCAGTTGGCGCATGGTGGCAAGTCCAAGGATGCTGGGAACAAGAGTATTGAAAGAGAAACAACTG
TCTCACCTGGAGGAGGCGGGAGGCCAGTTGGAAGGTTCTAGCTCTGGTTTAGAGGTACCCTAGGCTGGGT
CTTGTCTAAGGTATGGGTGGGAGAGAGGTGTAACCAGGTGGTGTCCAGTGTCCCAGCCCCAAGGCCCAAA
TCCTTACTCCTGATTCCTGCACTGAGGGGCAGCCTCTGCCAGTATCTCCTTTTGTGCTCAGCCGGGAGCT
AAGGCTTCCTGCCCTTTAGTCAGATGCTTTGAGACTGTGGGCTTCTGCCCTCCCGGCCCATGTCTCCCAC
CCCAACGGCACTCTTGCACATTCTGTTTCTCTCCCTTTCTCTCCACCTGACCATATCTGAGGGCGAACTT
AATTTTCCCAGAGGGAATTGCATTTGAGTCCAGTCACCTTACTGGGAGGCCATATCCTGTGGTTGAGCAG
GGAAATTATTTGTAGCCCTTCCAACATACTGACAGCGGTGGCTGATTTATCAGGACCTCTTGCCTGCTAA
TTTCATGCTTCAGTCCAATTCAGCTCCTTATTGAACTCTTCCTATTTTCCAGTAGCTGTTAGCAGCCTAA
GGACGGAGCTACCAATAAGACACAGTCCCCAGCCAAAGAGTGTCAGAAAATAATGACAATATGATGGGGT
GATGCTGTTATAGAGCTATTTATTTACCAAGTACTGCAAATACCCTGAGAAGGCATTAGTTGATCTGTCT
CCCTGGGAAGAAATCAGGAGGGTTTCAGAGGGGAAGTGTCATTTGAAAAGGGTTTTTGAGGGGTGAGTAG
GAGCTTACTATGCAAAGAAGCAGATGCACATTCCACATTGAGATCAAGAACGGTGGACAGTTACTTTATT
CATTTCTTCTTTCCTATCTATACGATTTGAGCTAGTTGGAAGGAAGCTGCTCAAACTTTAATGCGTATTT
TGTTAAAATGCAGATTCTGACTCAGTAGGTCTGGGGTGGGGCCCCAAGAGTTTGCAGTTTTGACAAGGAA
ACAGGTGGCGCCTATCAGATGCCCTGTAGCCAGCCATTGGCTACACTTTGAGCAGCAAAGCTTTAACATA
TGACCTTTTACCAACACTCACTGTTTGCCTTTACACGTGTGGCCCAGATTGGATGGAGGGCCTTGTCATT
CCTTGAAGGGTAGCGAAGGGCTGGTCGTTTGCAGGAAGGCCCTGGCTTGTCTTCTTCCAGAATGTGGTTA
TTCTCAGGCCCAGCCCAGTGAGTCCAGCACACCTACCAGATTGGAGCAGTGCTGTCAGAGAAACCCATTT
TCAACTCTGTAGCCCGCCCGTTTGGCTCTTCACCCAGACCTTGCACTACACGGGGTGATCACACATCTGT
GATTTTTAGGAAAATCACAGTTGTAAAACGATGGTCTTTTCTAAGAAGGCACCCTGTAGTGTATATTGAG
ACTCAATCTTTCTATAAGCATACATTCAAAAATCATTTTCTTAAGAAAGGGCCTTCCATAGTTTTGTTCC
CAGATGCCCGCACAGTCCTGTAGGAATGAGGGACTTACATCTGAATTCCTGGTACTCTGGGGAGGCCTTG
TGGCAAATGATTGGTTTCTCAAGTGCGGGCCTTTTGTGAAGTTGATGGGCTTGCTTTGACTCAGCCGATC
TTGGCCACATGTCTTTGAGTCATATGAAACGTGCCGGTAGCCTCAGGAACAGCACAGCCCGGAGAGGCGG
TGGCCAGCTGCCCGTCAGCTGTTCTGGGCTAATTTGGGGTTTCAGTGCTGGGGTGGAGAGAGTGCCGGGA
AATCCTAACAGGGGAGTTCTTGGGGCTTGAGGAAAGCCAGGCTGGGAAGGATGGAACGTGGACCTCTGCC

```
TCCCACCCCTCCATGAATCATGGTGCCAAGTAGAAATGCAGAGTTGATTCTCCCAGGGACAGAATTGCCC
AGCAGGGGATGTTGGCGTTTGGTGTGTGGCTGTTTCTGCTCATGTCCCCCGCTTGGCACACAAGAGACTG
CACTTGAAAACCACCCCTAGTCCCCAGCTCTTGTGCCCAGCAGCCCCTTCCCCAAATGCTTTACCCCCTG
TTCAATATTCTGCTGGTCTTTTAAGTCTCTCTTTCTCATCCTGAGTCTGGAAATCACTTGAGGGCAGGGA
CTCCACTTAGATCCTCGCACCTACTCAGTGCATGGCATGGAGGGGAGGCAGAATGAAAGGATGGACGTCT
GACCTCTTCCTCCTCCGAGAAATGCCGTAAGTGTGAGGCCAGCGGCCCCCGCGGAGTCCGGGTGGACAGC
CTCTCCTCTGCATCCTCCCTTCCTGGCAAGGATGGGAGAGAGTAAGTTTGCAGTCTGGCACACGCTCAGG
ACTCGATCGTGGGGCAAGATGTGATTTTTATGGCAGGAAAGTGGGAGACACATGATTAGAACTGGCTCTT
CCAGCGGAGACCCTAAGATCCTGCTCCTGCCCACAGTTGCAGCAACCTTCCCTTTTGGATTTCTAGGTCG
CTGGTAGAATAAGTTTTCCAGAATGGACTGTCTAGAGTTCCTTTCTTGCCAAGTGATTCAACCTGAGGGC
TCTGGGGTTTGATTTTTTTTGTTTTTTGTTTTTTTTTTTTAACTTTTGAGAAGTGTTGGGAAAATGGCTA
AGGAGAATTTCTTTAATTGTTCTGGGATTGGGAGTCTTCAAACTTGGATTTTAAAGAATTGCTATAGTTG
GTCATACCTCACATACATTAATATTAATATTAATACTAATATTAATACTAATGTTAATGATAATTACTTG
AATAAACACTTTGCCAGTCACAAAGCCCTTTTACACGATCTTTAATTTTTTTTCATTTTCTCAATAGCC
TTATGAGTGCCGCCTCCATGCTACAGATAAGGAAACTGAGGCTCAGGCAGTAGTTCACTCAGATGTAGTT
CAGAGAGTGATTAGGTGGGAGGGGGATCCAGGCCCCAGGACTTTGCATCTGGATCCTGAGTTCCTGGGTT
CTCCTCCCTGCCTCTCAGCCAGTGTTTCTGTGGGCCAGGGGGCGCCCACTGTGAACAGCTTCCTTTCCTG
AGGGACCCCGGAGAAGGTGCTCGCTGCATAGCAGACCTTTCTGAGTCAGGCCCATGAGGACTTGTAGGTC
CTCCTCTCTGCGCTGACATCATTGTGGCTGCATTGTGTGGAAATGCGGAGGGAGGCCAGTCTCACCGGGA
GCTTGGGAGTGGCCATGGCCCTTCAGAGAAGCCCGGATGAAGGCATGGGGCCTGGCCTTTGCACATTTGC
ACTAAGCAGTCATTGGGTGTGGTGGCAGCTCCTTTTGGCCAAAGGCAATGCCCAGAGAGGAGCTCAGCTG
TGAGCATTTGGCAGGCAGCTCTTGGGCAGCTAGAGGAATGGGTACTCAGGGGGCACCACAGCATCCACAG
CGTCAAAAATGGGGAATGGGCACTGCTTACTGTCAGTGGTGGACCACATTTGTTTTGGGAACAGGGAAGT
TGCTCTCTGTCCCTCAAATCATAACATTAGGCAAACTGCCTAACCAGCTTCTGTCTTGGCTTTGGGACTG
ACAGGATTCACTAATTGGAACCAACCTGACCTGTGAGGATGCTAACCTGTTTTTAAAAGGAATCAGTCTT
CATTACTTTTATGTATGAACACAGGGTGACTTGCTGTGTGCCCTTGGGGAAGTGACTCTCCTTCTCTGAG
TTTTACTTTCCTCATCTATAAAATGAGGATAAGGCTACTTTGAAAACTGTACTCTCCTGGGCGGGCTCGA
GCAGTGATTTTGATGGTCAGCATTGTCACATCCTCACATGGAAGTAACATTGTGAAATGTAAGGACAATA
TTTTCCATGCTTTTTTCTGTTCCAAGTCCTCTGTAATCCTGTGAAGCCTTTCTAGCCATCAGAAAGGGGA
ATGCTATGGCCTATACAAGATACAATAAGTGCAGAAAGCAGGTTGAAGTGGAATCCGTAGTCGCTCACCT
GCTCAGTAACCATCAGTGGCTCCCTATTGCCCATACAGTAAAGTGGGCAGACTCTGTCTGGATTTAAGAT
CCTCCTGTCTGGTTCCAAACCATCTTTCCAAACTCTTCTCTTGTGAAACTCCCCTTTTCCCTTCTTGCCT
CCCGGGAAGTCCCTGAGTGATCTTGGCTGTTCTCAACCATGGGCCTTTATTCCTGCTGCCTCAGCCCTTT
GAAATATGCTCCATAGTCCCTGCTTTGTGTTCCAAATAGATCTCCCCCTGATGGTAGAGCTTTGAAGTGC
TGTTCCTGGAAACTTAACG
```

**Figure S1.** Sequence of the oxytocin receptor gene targeted for the analysis. The areas marked in yellow are the CpG sites to be analyzed. Areas marked in gray indicate exons. The bases circled by squares indicate the translation start sites. The numbers of CpGs from 1 to 25 were assigned in order from upstream to downstream.

**Table S1.** List of papers published in this research project

| No | Bibliographic information of the article |
|---|---|
| 1 | Matsunaga M, Ohtsubo Y, Ishii K, Tsuboi H, Suzuki K, Takagishi H. Subjective well-being can be predicted by the caudate volume and promotion focus. *Brain Struct Funct* Online ahead of print (2024) |
| 2 | Kawamoto M, Takagishi H, Ishihara T, Takagi S, Kanai R, Sugihara G, Takahashi H, Matsuda T. Hippocampal volume mediates the relationship of parental rejection in childhood with social cognition in healthy adults. *Sci Rep* 2023;13:19167. |
| 3 | Tanaka H, Nishina N, Shou Q, Takahashi H, Sakagami M, Matsuda T, Inoue-Murayama M, Takagishi H. Association between arginine vasopressin receptor 1A (AVPR1A) polymorphism and inequity aversion. *Proc Biol Sci*. 2023;290(2000):20230378. |
| 4 | Tanaka H, Shou Q, Kiyonari T, Matsuda T, Sakagami M, Takagishi H. Right dorsolateral prefrontal cortex regulates default prosociality preference. *Cereb Cortex.* 2023;33:5420–5425. |
| 5 | Fermin ASR, et al. The neuroanatomy of social trust predicts depression vulnerability. *Sci Rep* 2022;12:16724. |
| 6 | Shou Q, Yamada J, Nishina K, Matsunaga M, Matsuda T, Takagishi H. Association between salivary oxytocin levels and the amygdala and hippocampal volumes. *Brain Struct Funct.* 2022;227:2503–2511. |
| 7 | Shou Q, Yamada J, Nishina K, Matsunaga M, Kiyonari T, Takagishi H. Is oxytocin a trust hormone? Salivary oxytocin is associated with caution but not with general trust. *PLoS On*. 2022;17:e0267988. |
| 8 | Nishina K, Shou Q, Takahashi H, Sakagami M, Inoue-Murayama M, Takagishi H. Association between polymorphism (5-HTTLPR) of the serotonin transporter gene and behavioral response to unfair distribution. *Front Behav Neurosci.* 2022;16:762092. |
| 9 | Yamada J, Nakawake Y, Shou Q, Nishina K, Matsunaga M, Takagishi H. Salivary oxytocin is negatively associated with religious faith in Japanese non-Abrahamic people. *Front Psychol.* 2012:12;705781. |
| 10 | Ishihara T, Miyazaki A, Tanaka H, Fujii T, Takahashi M, Nishina K, Kanari K, Takagishi H, Matsuda T. Childhood exercise predicts response inhibition in later life via changes in brain connectivity and structure. *NeuroImage.* 2021;237:118196. |
| 11 | Nishina K, Takagishi H, Takahashi H, Sakagami M, Inoue-Murayama M. Association of polymorphism of arginine-vasopressin receptor 1A (AVPR1a) gene with trust and reciprocity. *Front Hum Neurosci.* 2019;13:230. |
| 12 | Nishina K, Takagishi H, Fermin ASR, Inoue-Murayama M, Takahashi H, Sakagami M, Yamagishi T. Association of the oxytocin receptor gene with attitudinal trust: Role of amygdala volume. *Cogn Affect Neurosci.* 2018;13:1091–1097. |

13 Yamagishi T, Li Y, Fermin ASR, Kanai R, Takagishi H, Matsumoto Y, Kiyonari T, Sakagami M. Behavioural differences and neural substrates of altruistic and spiteful punishment. *Sci Rep.* 2017;7:14654.

14 Yamagishi T, Matsumoto Y, Kiyonari T, Takagishi H, Li Y, Kanai R, Sakagami M. Response time in economic games reflects different types of decision conflict for prosocial and proself individuals. *Proc Natl Acad Sci USA.* 2017;114:6394–6399.

15 Yamagishi T, Takagishi H, Fermin ASR, Kanai R, Li Y, Matsumoto Y. Cortical thickness of the dorsolateral prefrontal cortex predicts strategic choices in economic games. *Proc Natl Acad Sci USA.* 2016:113:5582–5587.

16 Matsumoto Y, Yamagishi T, Li Y, Kiyonari T. Prosocial behavior increases with age across five economic games. *PLoS One.* 2016;11:e0158671.

17 Yamagishi T, Li Y, Matsumoto Y, Kiyonari T. Moral bargain hunters purchase moral righteousness when it is cheap: within-individual effect of stake size in economic games. *Sci Rep.* 2016;6:27824.

18 Nishina K, Takagishi H, Inoue-Murayama M, Takahashi H, Yamagishi T. Polymorphism of the oxytocin receptor gene modulates behavioral and attitudinal trust among men but not women. *PLoS One.* 2015;10;e0137089.

19 Yamagishi T, Li Y, Takagishi H, Matsumoto Y, Kiyonari T. In search of homo economicus. *Psychol Sci.* 2014;25:1699–1711.

**Table S2** Effect of relational mobility on methylation at CpG 1

| | β | SE | t | p | CI | | df |
|---|---|---|---|---|---|---|---|
| | | | | | Lower | Upper | |
| (Intercept) | .000 | .095 | | | | | |
| Age | .487 | .093 | 5.231 | < .0001 | .302 | .672 | 90 |
| Sex | .045 | .094 | 0.479 | .633 | -.141 | .231 | 90 |
| % of epithelial cells | .084 | .134 | 0.627 | .533 | -.182 | .349 | 90 |
| Relational Mobility | .046 | .095 | 0.488 | .627 | -.142 | .234 | 90 |
| Adjusted $R^2$ | .199 | | | | | | |

*β: standardized partial regression coefficient, SE: standard error, CI: confidence interval

**Table S3** Effect of relational mobility on methylation at CpG 2

| | β | SE | t | P | CI | | df |
|---|---|---|---|---|---|---|---|
| | | | | | Lower | Upper | |
| (Intercept) | .000 | .103 | | | | | |
| Age | .045 | .094 | 0.482 | .631 | -.142 | .232 | 90 |
| Sex | -.137 | .105 | -1.305 | .195 | -.347 | .072 | 90 |
| % of epithelial cells | -.210 | .106 | -1.992 | .049 | -.420 | -.001 | 90 |
| Relational Mobility | .005 | .069 | 0.080 | .937 | -.131 | .142 | 90 |
| Adjusted $R^2$ | .035 | | | | | | |

*β: standardized partial regression coefficient, SE: standard error, CI: confidence interval

**Table S4** Effect of relational mobility on methylation at CpG 3

| | β | SE | t | p | CI | | df |
|---|---|---|---|---|---|---|---|
| | | | | | Lower | Upper | |
| (Intercept) | .000 | .101 | | | | | |
| Age | .330 | .096 | 3.439 | .0009 | .139 | .520 | 90 |
| Sex | -.186 | .099 | -1.887 | .062 | -.382 | .010 | 90 |
| % of epithelial cells | .136 | .137 | 0.992 | .324 | -.136 | .408 | 90 |
| Relational Mobility | .093 | .118 | 0.786 | .434 | -.142 | .328 | 90 |
| Adjusted $R^2$ | .108 | | | | | | |

*β: standardized partial regression coefficient, SE: standard error, CI: confidence interval

**Table S5** Effect of relational mobility on methylation at CpG 4

| | | | | | CI | | |
|---|---|---|---|---|---|---|---|
| | **β** | **SE** | **t** | **p** | **Lower** | **Upper** | **df** |
| (Intercept) | .000 | .105 | | | | | |
| Age | -.019 | .111 | -0.169 | .867 | -.239 | .202 | 90 |
| Sex | -.192 | .103 | -1.858 | .066 | -.397 | .013 | 90 |
| % of epithelial cells | .013 | .122 | 0.105 | .917 | -.229 | .254 | 90 |
| Relational Mobility | -.181 | .111 | -1.634 | .106 | -.401 | .039 | 90 |
| Adjusted $R^2$ | .018 | | | | | | |

*β: standardized partial regression coefficient, SE: standard error, CI: confidence interval

**Table S6** Effect of relational mobility on methylation at CpG 5

| | | | | | CI | | |
|---|---|---|---|---|---|---|---|
| | **β** | **SE** | **t** | **p** | **Lower** | **Upper** | **df** |
| (Intercept) | .000 | .087 | | | | | |
| Age | .134 | .068 | 1.974 | .051 | -.001 | .268 | 90 |
| Sex | -.171 | .080 | -2.134 | .036 | -.329 | -.012 | 90 |
| % of epithelial cells | -.502 | .108 | -4.661 | <.0001 | -.716 | -.288 | 90 |
| Relational Mobility | .042 | .074 | 0.573 | .568 | -.104 | .188 | 90 |
| Adjusted $R^2$ | .318 | | | | | | |

*β: standardized partial regression coefficient, SE: standard error, CI: confidence interval

**Table S7** Effect of relational mobility on methylation at CpG 6

| | | | | | CI | | |
|---|---|---|---|---|---|---|---|
| | **β** | **SE** | **t** | **p** | **Lower** | **Upper** | **df** |
| (Intercept) | .000 | .097 | | | | | |
| Age | .379 | .102 | 3.709 | .0004 | .176 | .582 | 90 |
| Sex | -.130 | .096 | -1.355 | .179 | -.321 | .061 | 90 |
| % of epithelial cells | .275 | .105 | 2.621 | .010 | .067 | .483 | 90 |
| Relational Mobility | .099 | .111 | 0.892 | .375 | -.122 | .321 | 90 |
| Adjusted $R^2$ | .164 | | | | | | |

*β: standardized partial regression coefficient, SE: standard error, CI: confidence interval

**Table S8** Effect of relational mobility on methylation at CpG 7

| | | | | | CI | | |
|---|---|---|---|---|---|---|---|
| | **β** | **SE** | **t** | **p** | **Lower** | **Upper** | **df** |
| (Intercept) | .000 | .102 | | | | | |
| Age | .190 | .098 | 1.935 | .056 | -.005 | .385 | 90 |
| Sex | -.074 | .099 | -0.741 | .461 | -.271 | .124 | 90 |
| % of epithelial cells | -.217 | .126 | -1.725 | .088 | -.467 | .033 | 90 |
| Relational Mobility | .047 | .118 | 0.399 | .691 | -.187 | .281 | 90 |
| Adjusted $R^2$ | .065 | | | | | | |

*β: standardized partial regression coefficient, SE: standard error, CI: confidence interval

**Table S9** Effect of relational mobility on methylation at CpG 8

| | | | | | CI | | |
|---|---|---|---|---|---|---|---|
| | **β** | **SE** | **t** | **p** | **Lower** | **Upper** | **df** |
| (Intercept) | .000 | .105 | | | | | |
| Age | .107 | .116 | 0.926 | .357 | -.123 | .337 | 90 |
| Sex | -.116 | .106 | -1.095 | .277 | -.326 | .094 | 90 |
| % of epithelial cells | .162 | .096 | 1.681 | .096 | -.030 | .353 | 90 |
| Relational Mobility | .093 | .121 | 0.770 | .443 | -.147 | .333 | 90 |
| Adjusted $R^2$ | .003 | | | | | | |

*β: standardized partial regression coefficient, SE: standard error, CI: confidence interval

**Table S10** Effect of relational mobility on methylation at CpG 9

| | | | | | CI | | |
|---|---|---|---|---|---|---|---|
| | **β** | **SE** | **t** | **p** | **Lower** | **Upper** | **df** |
| (Intercept) | .000 | .105 | | | | | |
| Age | .055 | .104 | 0.529 | .598 | -.152 | .262 | 90 |
| Sex | -.195 | .109 | -1.780 | .078 | -.412 | .023 | 90 |
| % of epithelial cells | .175 | .136 | 1.289 | .201 | -.095 | .444 | 90 |
| Relational Mobility | .121 | .108 | 1.120 | .266 | -.093 | .334 | 90 |
| Adjusted $R^2$ | .028 | | | | | | |

*β: standardized partial regression coefficient, SE: standard error, CI: confidence interval

**Table S11** Effect of relational mobility on methylation at CpG 10

| | | | | | CI | | |
|---|---|---|---|---|---|---|---|
| | **β** | **SE** | **t** | **p** | **Lower** | **Upper** | **df** |
| (Intercept) | .000 | .089 | | | | | |
| Age | .065 | .075 | 0.862 | .391 | -.085 | .215 | 90 |
| Sex | -.135 | .084 | -1.606 | .112 | -.303 | .032 | 90 |
| % of epithelial cells | -.518 | .117 | -4.428 | <.0001 | -.751 | -.286 | 90 |
| Relational Mobility | -.023 | .084 | -0.267 | .790 | -.190 | .145 | 90 |
| Adjusted $R^2$ | .284 | | | | | | |

*β: standardized partial regression coefficient, SE: standard error, CI: confidence interval

**Table S12** Effect of relational mobility on methylation at CpG 11

| | | | | | CI | | |
|---|---|---|---|---|---|---|---|
| | **β** | **SE** | **t** | **p** | **Lower** | **Upper** | **df** |
| (Intercept) | .000 | .097 | | | | | |
| Age | -.094 | .100 | -0.947 | .346 | -.292 | .104 | 90 |
| Sex | .256 | .096 | 2.665 | .009 | .065 | .447 | 90 |
| % of epithelial cells | .001 | .109 | 0.005 | .996 | -.216 | .217 | 90 |
| Relational Mobility | -.325 | .095 | -3.438 | .0009 | -.513 | -.137 | 90 |
| Adjusted $R^2$ | .157 | | | | | | |

*β: standardized partial regression coefficient, SE: standard error, CI: confidence interval

**Table S13** Effect of relational mobility on methylation at CpG 12

| | | | | | CI | | |
|---|---|---|---|---|---|---|---|
| | **β** | **SE** | **t** | **p** | **Lower** | **Upper** | **df** |
| (Intercept) | .000 | .104 | | | | | |
| Age | -.052 | .109 | -0.477 | .634 | -.268 | .165 | 90 |
| Sex | -.178 | .106 | -1.680 | .096 | -.389 | .033 | 90 |
| % of epithelial cells | .180 | .114 | 1.579 | .118 | -.046 | .406 | 90 |
| Relational Mobility | .167 | .133 | 1.258 | .212 | -.097 | .430 | 90 |
| Adjusted $R^2$ | .044 | | | | | | |

*β: standardized partial regression coefficient, SE: standard error, CI: confidence interval

**Table S14** Effect of relational mobility on methylation at CpG 13

| | β | SE | t | p | CI | | df |
|---|---|---|---|---|---|---|---|
| | | | | | Lower | Upper | |
| (Intercept) | .000 | .104 | | | | | |
| Age | -.009 | .115 | -0.082 | .935 | -.237 | .218 | 90 |
| Sex | -.175 | .108 | -1.629 | .107 | -.389 | .039 | 90 |
| % of epithelial cells | .222 | .101 | 2.205 | .030 | .022 | .422 | 90 |
| Relational Mobility | -.023 | .111 | -0.211 | .833 | -.244 | .197 | 90 |
| Adjusted $R^2$ | .028 | | | | | | |

*β: standardized partial regression coefficient, SE: standard error, CI: confidence interval

**Table S15** Effect of relational mobility on methylation at CpG 14

| | β | SE | t | p | CI | | df |
|---|---|---|---|---|---|---|---|
| | | | | | Lower | Upper | |
| (Intercept) | .000 | .099 | | | | | |
| Age | .032 | .099 | 0.326 | .745 | -.164 | .228 | 90 |
| Sex | -.218 | .092 | -2.387 | .019 | -.400 | -.037 | 90 |
| % of epithelial cells | .396 | .137 | 2.891 | .005 | .124 | .669 | 90 |
| Relational Mobility | -.043 | .113 | -0.379 | .705 | -.267 | .181 | 90 |
| Adjusted $R^2$ | .147 | | | | | | |

*β: standardized partial regression coefficient, SE: standard error, CI: confidence interval

**Table S16** Effect of relational mobility on methylation at CpG 15

| | β | SE | t | P | CI | | df |
|---|---|---|---|---|---|---|---|
| | | | | | Lower | Upper | |
| (Intercept) | .000 | .097 | | | | | |
| Age | -.103 | .094 | -1.103 | .273 | -.290 | .083 | 90 |
| Sex | -.179 | .097 | -1.839 | .069 | -.372 | .014 | 90 |
| % of epithelial cells | -.385 | .114 | -3.389 | .001 | -.611 | -.159 | 90 |
| Relational Mobility | -.054 | .114 | -0.477 | .635 | -.281 | .172 | 90 |
| Adjusted $R^2$ | .160 | | | | | | |

*β: standardized partial regression coefficient, SE: standard error, CI: confidence interval

**Table S17** Effect of relational mobility on methylation at CpG 16

| | β | SE | t | p | CI | | df |
|---|---|---|---|---|---|---|---|
| | | | | | Lower | Upper | |
| (Intercept) | .000 | .105 | | | | | |
| Age | .058 | .101 | 0.578 | .564 | -.142 | .259 | 90 |
| Sex | -.162 | .116 | -1.399 | .165 | -.391 | .068 | 90 |
| % of epithelial cells | -.095 | .115 | -0.831 | .408 | -.323 | .133 | 90 |
| Relational Mobility | -.050 | .096 | -0.517 | .606 | -.241 | .142 | 90 |
| Adjusted $R^2$ | .001 | | | | | | |

*β: standardized partial regression coefficient, SE: standard error, CI: confidence interval

**Table S18** Effect of relational mobility on methylation at CpG 17

| | β | SE | t | p | CI | | df |
|---|---|---|---|---|---|---|---|
| | | | | | Lower | Upper | |
| (Intercept) | .000 | .106 | | | | | |
| Age | -.050 | .098 | -0.508 | .613 | -.243 | .144 | 90 |
| Sex | -.077 | .099 | -0.783 | .436 | -.273 | .119 | 90 |
| % of epithelial cells | -.127 | .153 | -0.832 | .408 | -.431 | .176 | 90 |
| Relational Mobility | -.189 | .113 | -1.673 | .098 | -.414 | .035 | 90 |
| Adjusted $R^2$ | .004 | | | | | | |

*β: standardized partial regression coefficient, SE: standard error, CI: confidence interval

**Table S19** Effect of relational mobility on methylation at CpG 18

| | β | SE | t | p | CI | | df |
|---|---|---|---|---|---|---|---|
| | | | | | Lower | Upper | |
| (Intercept) | .000 | .098 | | | | | |
| Age | .049 | .100 | 0.484 | .630 | -.151 | .248 | 90 |
| Sex | .013 | .099 | 0.129 | .898 | -.184 | .209 | 90 |
| % of epithelial cells | -.404 | .102 | -3.942 | .0002 | -.607 | -.200 | 90 |
| Relational Mobility | .028 | .113 | 0.245 | .807 | -.197 | .253 | 90 |
| Adjusted $R^2$ | .135 | | | | | | |

*β: standardized partial regression coefficient, SE: standard error, CI: confidence interval

**Table S20** Effect of relational mobility on methylation at CpG 19

| | | | | | CI | | |
|---|---|---|---|---|---|---|---|
| | β | SE | t | p | Lower | Upper | df |
| (Intercept) | .000 | .096 | | | | | |
| Age | -.109 | .095 | -1.149 | .254 | -.297 | .079 | 90 |
| Sex | -.159 | .095 | -1.675 | .098 | -.348 | .030 | 90 |
| % of epithelial cells | -.231 | .105 | -2.205 | .030 | -.439 | -.023 | 90 |
| Relational Mobility | .270 | .109 | 2.476 | .015 | .053 | .486 | 90 |
| Adjusted $R^2$ | .180 | | | | | | |

*β: standardized partial regression coefficient, SE: standard error, CI: confidence interval

**Table S21** Effect of relational mobility on methylation at CpG 20

| | | | | | CI | | |
|---|---|---|---|---|---|---|---|
| | β | SE | t | p | Lower | Upper | df |
| (Intercept) | .000 | .100 | | | | | |
| Age | -.050 | .099 | -0.506 | .614 | -.246 | .146 | 90 |
| Sex | -.187 | .101 | -1.841 | .069 | -.388 | .015 | 90 |
| % of epithelial cells | .049 | .127 | 0.383 | .703 | -.204 | .302 | 90 |
| Relational Mobility | .298 | .111 | 2.676 | .009 | .077 | .519 | 90 |
| Adjusted $R^2$ | .101 | | | | | | |

*β: standardized partial regression coefficient, SE: standard error, CI: confidence interval

**Table S22** Effect of relational mobility on methylation at CpG 21

| | | | | | CI | | |
|---|---|---|---|---|---|---|---|
| | β | SE | t | p | Lower | Upper | df |
| (Intercept) | .000 | .082 | | | | | |
| Age | .100 | .073 | 1.362 | .177 | -.046 | .246 | 90 |
| Sex | -.191 | .078 | -2.467 | .016 | -.346 | -.037 | 90 |
| % of epithelial cells | -.573 | .106 | -5.428 | <.0001 | -.783 | -.364 | 90 |
| Relational Mobility | -.001 | .077 | -0.014 | .989 | -.155 | .153 | 90 |
| Adjusted $R^2$ | .393 | | | | | | |

*β: standardized partial regression coefficient, SE: standard error, CI: confidence interval

**Table S23** Effect of relational mobility on methylation at CpG 22

| | | | | | CI | | |
|---|---|---|---|---|---|---|---|
| | **β** | **SE** | **t** | **p** | **Lower** | **Upper** | **df** |
| (Intercept) | .000 | .102 | | | | | |
| Age | -.012 | .096 | -0.129 | .897 | -.204 | .179 | 90 |
| Sex | -.238 | .102 | -2.328 | .022 | -.441 | -.035 | 90 |
| % of epithelial cells | .247 | .116 | 2.128 | .036 | .016 | .478 | 90 |
| Relational Mobility | -.074 | .119 | -0.624 | .534 | -.310 | .162 | 90 |
| Adjusted $R^2$ | .068 | | | | | | |

*β: standardized partial regression coefficient, SE: standard error, CI: confidence interval

**Table S24** Effect of relational mobility on methylation at CpG 23

| | | | | | CI | | |
|---|---|---|---|---|---|---|---|
| | **β** | **SE** | **t** | **p** | **Lower** | **Upper** | **df** |
| (Intercept) | .000 | .104 | | | | | |
| Age | .264 | .103 | 2.552 | .012 | .058 | .469 | 90 |
| Sex | -.137 | .100 | -1.374 | .173 | -.335 | .061 | 90 |
| % of epithelial cells | .001 | .127 | 0.005 | .996 | -.252 | .253 | 90 |
| Relational Mobility | .061 | .120 | 0.505 | .615 | -.178 | .299 | 90 |
| Adjusted $R^2$ | .049 | | | | | | |

*β: standardized partial regression coefficient, SE: standard error, CI: confidence interval

**Table S25** Effect of relational mobility on methylation at CpG 24

| | | | | | CI | | |
|---|---|---|---|---|---|---|---|
| | **β** | **SE** | **t** | **p** | **Lower** | **Upper** | **df** |
| (Intercept) | .000 | .099 | | | | | |
| Age | .140 | .099 | 1.414 | .161 | -.057 | .336 | 90 |
| Sex | .256 | .103 | 2.480 | .015 | .051 | .462 | 90 |
| % of epithelial cells | .111 | .115 | 0.967 | .336 | -.118 | .340 | 90 |
| Relational Mobility | -.181 | .105 | -1.726 | .088 | -.390 | .027 | 90 |
| Adjusted $R^2$ | .128 | | | | | | |

*β: standardized partial regression coefficient, SE: standard error, CI: confidence interval

**Table S26** Effect of relational mobility on methylation at CpG 25

| | β | SE | t | p | CI Lower | CI Upper | df |
|---|---|---|---|---|---|---|---|
| (Intercept) | .000 | .057 | | | | | |
| Age | -.047 | .059 | -0.791 | .431 | -.163 | .070 | 90 |
| Sex | .013 | .059 | 0.215 | .831 | -.104 | .129 | 90 |
| % of epithelial cells | -.865 | .058 | -14.901 | <.0001 | -.981 | -.750 | 90 |
| Relational Mobility | -.094 | .061 | -1.549 | .125 | -.215 | .027 | 90 |
| Adjusted $R^2$ | .705 | | | | | | |

*β: standardized partial regression coefficient, SE: standard error, CI: confidence interval
